# Knockout of a single *Sox* gene resurrects an ancestral cell type in the sea anemone *Nematostella vectensis*

**DOI:** 10.1101/2021.09.30.462561

**Authors:** Leslie S. Babonis, Camille Enjolras, Abigail J. Reft, Brent M. Foster, Fredrik Hugosson, Joseph F. Ryan, Marymegan Daly, Mark Q. Martindale

## Abstract

Cnidocytes are the explosive stinging cells found only in cnidarians (corals, jellyfish, etc). Specialized for prey capture and defense, cnidocytes are morphologically complex and vary widely in form and function across taxa; how such diversity evolved is unknown. Using CRISPR/Cas9-mediated genome editing in the burrowing sea anemone *Nematostella vectensis*, we show that a single transcription factor (*NvSox2*) acts as a binary switch between two alternative cnidocyte fates. Knockout of *NvSox2* caused a complete transformation of nematocytes (piercing cells) into spirocytes (ensnaring cells). The type of spirocyte induced by *NvSox2* knockout (robust spirocyte) is not normally found in *N. vectensis* but is common in sea anemones from other habitats. Homeotic control of cell fate provides a mechanistic explanation for the discontinuous distribution of cnidocyte types across cnidarians and demonstrates how simple counts of cell types can underestimate biodiversity.

## Main

Novel cell types arise frequently and are found in many animal lineages; yet understanding the mechanisms that underly the origin of new cell phenotypes is a persistent challenge in biology (*1, 2*). Homeosis, the transformation of one part of an organism into another part (*3*), is a well-studied driver of morphological evolution in multicellular organisms (*4*) and often presents as the conversion of one serial segment into another following the manipulation of regionally expressed regulatory genes (*5, 6*). Studies of neural subtype differentiation nematodes, flies, and vertebrates have also identified single genetic switches sufficient to transform one cell type into another, suggesting homeosis may also promote the evolution of novelty at the level of individual cells (*7*). To date, however, no study has functionally demonstrated homeotic induction of atavism – the reemergence of an ancestral character – at the level of an individual cell type. Such a demonstration would provide a mechanistic understanding of how homeosis could drive cell type diversification.

Cnidocytes (stinging cells) are among the most morphologically complex of all cell types and are an iconic example of a cellular novelty (**Fig 1**). Found exclusively in cnidarians (corals, jellyfish, and their relatives), cnidocytes comprise a diverse lineage of cells specialized for prey capture, defense, and habitat use that vary widely in both form and function across taxa (*8*). Indeed, the “cnidom”, or inventory of cnidocyte types found in each species, is an important diagnostic character for understanding cnidarian taxonomy (*9*). Three major subtypes of cnidocyte are currently recognized (*10*). The first, nematocytes (**Fig 1A-C**), have an explosive organelle typically containing a spiny piercing apparatus and toxic venom (*11*). Across cnidarians, over 30 different types of nematocytes have been described, differing largely in the morphology of the eversible tubule (*8*). The second cnidocyte subtype, spirocytes, exhibit much less morphological variation and are described only as gracile or robust (*12*). These cells are typified by a thin capsule wall and an eversible tubule adorned with fine lateral rods to create an ensnaring web (*13*) (**Fig 1D-F**). Ptychocytes vary in size but have no other described differences and their payload consists of only a sticky tubule (*14*) (**Fig 1G-I**). The phylogenetic distribution of cnidocyte subtypes across cnidarians suggests that spirocytes and ptychocytes evolved from a nematocyte ancestor (**Fig 1J**); however, the regulatory mechanisms driving cnidocyte diversification have not been characterized so the orthology among cnidocyte subtypes remains unknown.

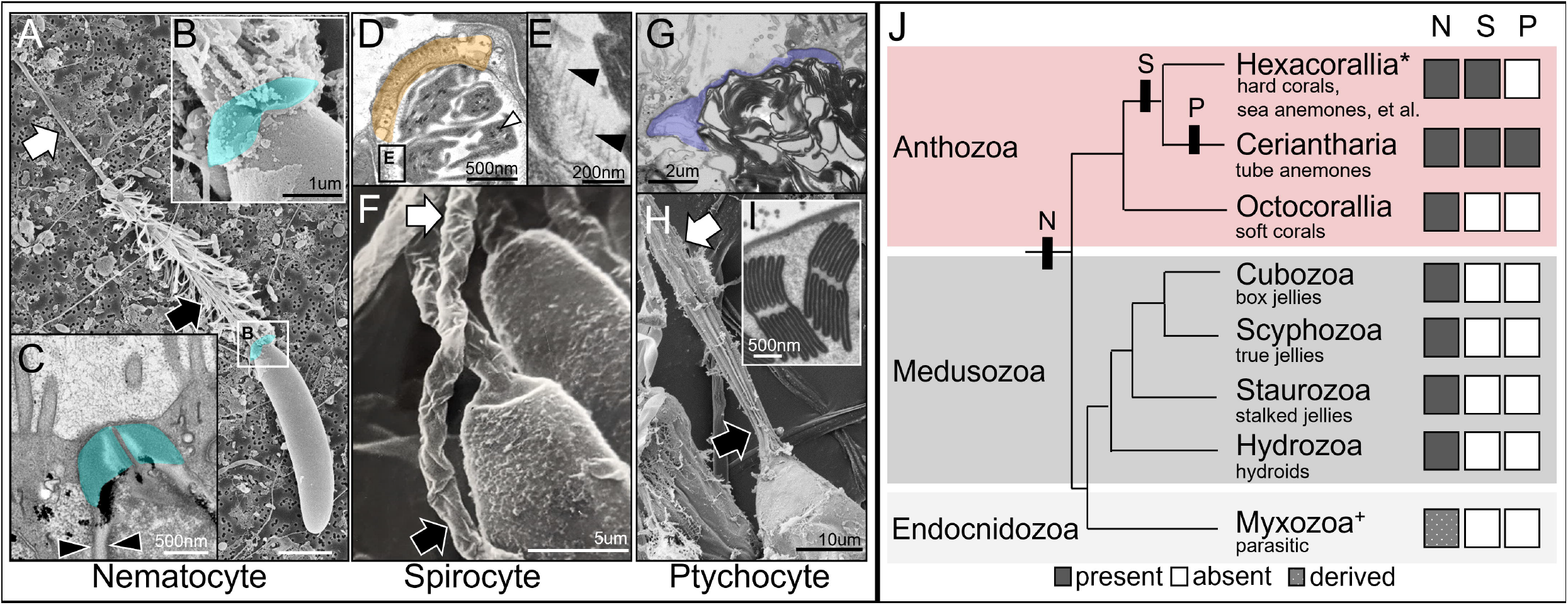

Sox genes play an inductive role in cell fate decisions across animals (*15*). Tissue-restricted expression of various Sox orthologs has been demonstrated previously in cnidarians and is consistent with a role for these transcription factors in patterning distinct cell types (*16–20*). Using the starlet sea anemone, *Nematostella vectensis*, we demonstrate that a single Sox gene, *NvSox2*, acts as a homeotic selector regulating the switch between two alternative cnidocyte fates: nematocyte or spirocyte. Thus, we present functional evidence of homeotic control of cell fate in a cnidarian and provide a framework for understanding the discontinuous variation in cnidocyte identity observed across cnidarians.

### Knockout of NvSox2 transforms nematocytes into spirocytes

*NvSox2* is one of 14 Sox genes in *N. vectensis*, only two of which (*NvSoxB2* and *NvSox2*) are expressed in a pattern consistent with specification of cell identity during early embryogenesis (*16*). We examined the expression of *NvSox2* further and found it to be co-expressed with the cnidocyte-specific transcription factor *PaxA* in a subset of developing cnidocytes (**Fig 2A**) early in development. *NvSox2* expression is suppressed following knockdown of *NvSoxB2* (expressed in progenitors of neurons and cnidocytes) (*18*) but is not affected in response to knockdown of *PaxA* (**Fig 2B**), suggesting NvSox2 is downstream of *NvSoxB2* and upstream of *PaxA* in developing cnidocytes (**Fig 2C**). To understand the function of *NvSox2* in cnidocyte specification, we used CRISPR/Cas9-mediated genome editing to generate F2 homozygous knockouts for the *NvSox2* locus (**Supplementary Fig S1, Supplementary Data S1**). *NvSox2* is first expressed at the blastula stage and this expression is completely abolished in *NvSox2* mutants (**Fig 2D**, **Supplementary Fig S2**). We allowed these *Sox2* mutants to grow to the juvenile polyp stage to investigate the morphological effects of *NvSox2* knockout.

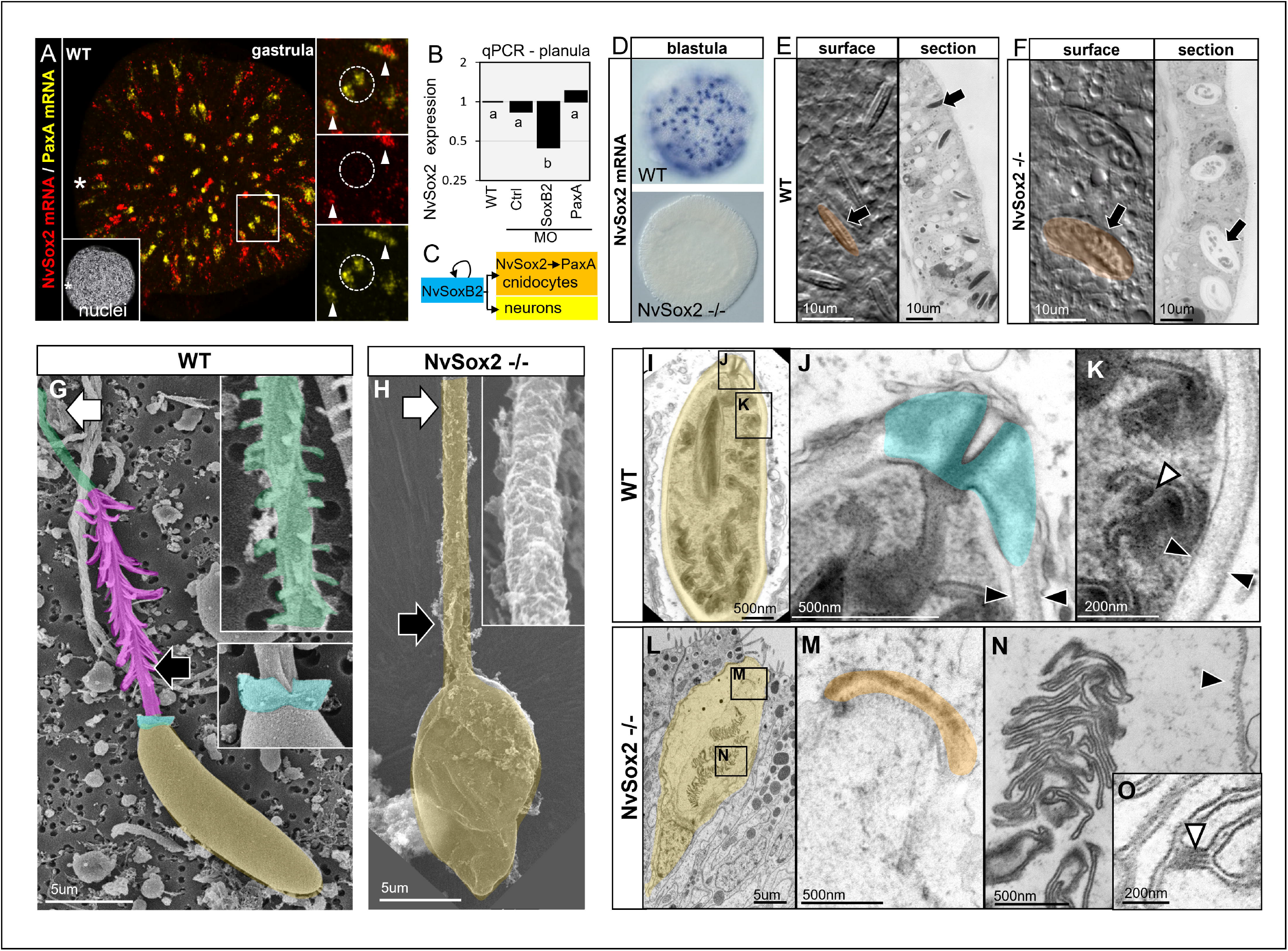

At the polyp stage, the body wall of *N. vectensis* is populated almost exclusively by a single type of nematocyte, the small basitrichous isorhiza (*21, 22*). In recently metamorphosed polyps (4-tentacle stage), we found the body wall nematocytes in *NvSox2* mutants to be completely replaced by a morphologically distinct cell type (**Fig 2E,F**). At a gross level, the mutant cell type had the appearance of a robust spirocyte, a type of spirocyte not normally found in *N. vectensis* but common in other sea anemones (**Supplementary Data S2**). To investigate this further, we examined the morphology of these cells using scanning and transmission electron microscopy (SEM and TEM, respectively).

Discharged nematocytes from wildtype (WT) polyps had rigid capsules with prominent apical flaps and an extrusible tubule with large spines proximally and small spines distally (**Fig 2G**). These features are characteristic of sea anemone nematocytes (*23–25*). By contrast, the capsule of the discharged mutant cells was thin and collapsed upon discharge; further, this capsule lacked apical flaps and the tubule discharged from these mutant cells was smooth along its entire length (**Fig 2H**). These features are consistent with spirocytes, but not nematocytes (*13, 26, 27*). Examination of the mutant cnidocytes using TEM (**Fig 2I-O**) further support these observations. Unlike nematocytes which have a thick capsule wall extending away from the sealed apical flaps (**Fig 2I-K**), the mutant cells have a thin capsule wall with a serrated appearance along the inner surface and a flat apical plate/cap (**Fig 2L-N**). Furthermore, the internalized/coiled tubule exhibits small rods consistent with the appearance of undischarged spirocytes (**Fig 2O**), although these rods were smaller and few in number than has been described for other spirocytes (*13*). Taken together, these results demonstrate that knockout of *NvSox2* in *N. vectensis* causes a transformation in cnidocytes from a piercing phenotype typical of nematocytes into an ensnaring phenotype typical of robust spirocytes.

### Assembly of the NvSox2-mediated nematocyte gene regulatory network

To understand the molecular regulation of this shift in cnidocyte phenotype, we examined the expression of genes known to be specific to different subtypes of cnidocytes in *N. vectensis* (*25, 28*). Expression of nematocyte-specific transcription factor *PaxA* is completely abolished in the body wall of *NvSox2* mutants (**Fig 3A; Supplementary Fig S3**). Likewise, expression of the nematocyte-specific molecule minicollagen 1 (*Mcol1*) was also abolished in the body wall of tentacle bud stage mutants, while *Mcol4*, which is expressed in both nematocytes and spirocytes, was unaffected (**Fig 3B**). Quantitative analysis of cnidocyte development shows that 80% of the Mcol4-labeled developing cnidocytes in the body wall of WT animals also express *Mcol1* and this proportion is significantly reduced to less than 10% in *NvSox2* mutants (**Fig 3C**) (Mann Whitney U; p <0.01). These results confirm that knockout of *NvSox2* did not affect the total number of cnidocytes specified in the body wall but transformed the identity of the cnidocytes from nematocyte to spirocyte.

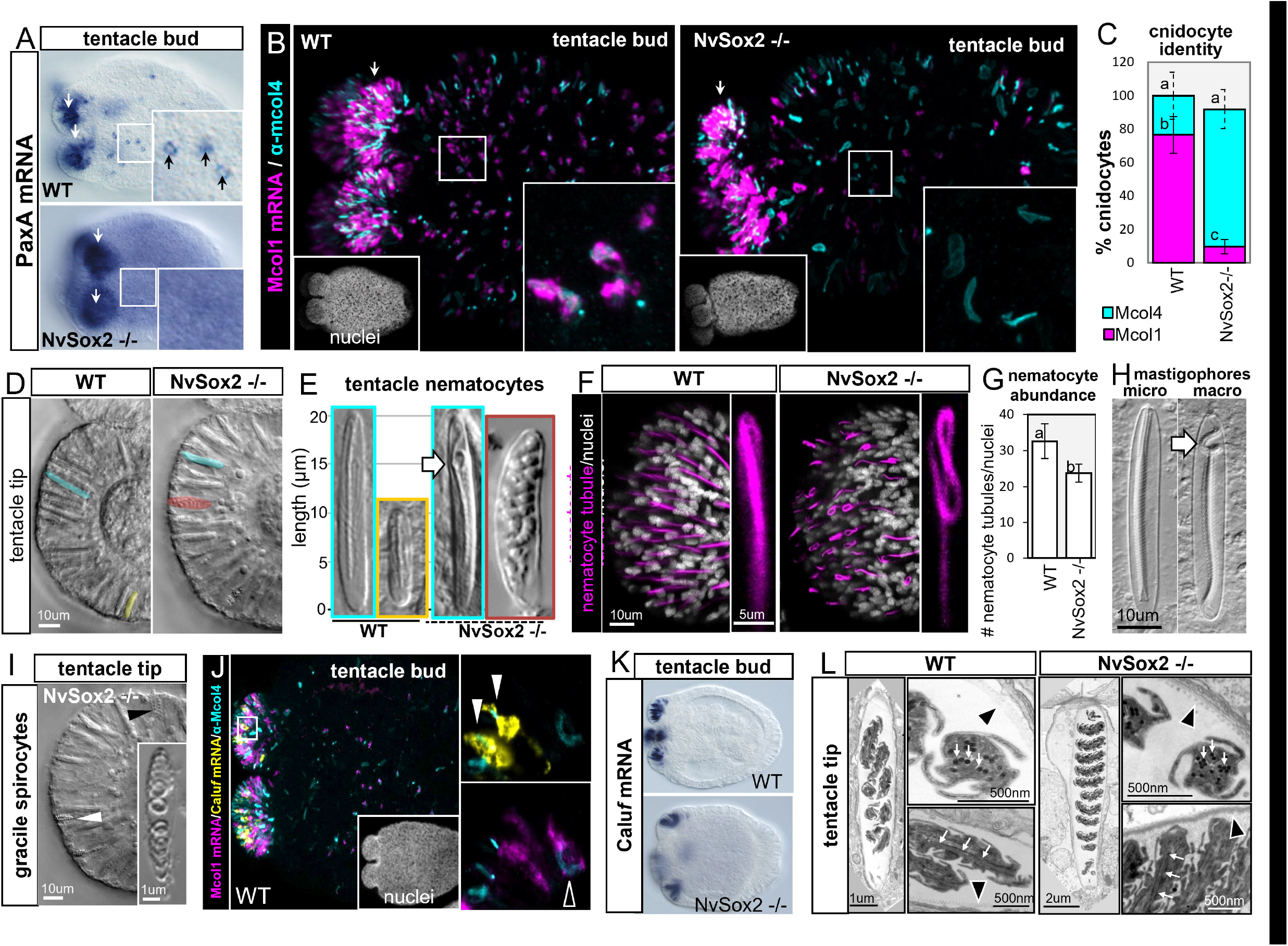

### NvSox2 is not required for specification of other types of cnidocytes

The expression of *PaxA* and *Mcol1* did not appear to be affected in the tentacle tips of *NvSox2* mutants (**Fig 3A-C**); however, examination of the tentacle tips by light microscopy confirmed that the small basitrichous isorhizas were completely transformed into robust spirocytes, mirroring the effects in the body wall (**Fig 3D-E**). The tentacle tips of *N. vectensis* are populated largely by spirocytes and large basitrichous isorhiza nematocytes (*22, 25, 28*). Further examination of the large basitrichous isorhizas using H_2_O_2_-conjugated tyramide revealed normal abundance and distribution of this cell type but aberrant tubule morphology within the stinging capsule (**Fig 3F,G**). Specifically, the proximal portion of the tubule appeared looped in *NvSox2* mutants, as compared to the straight tubule found in large basitrichous isorhizas from WT polyps. Induced discharge of these cells in both WT and mutant polyps using an infrared laser ablation system (XY Clone, Hamilton Thorne, USA) confirmed that the looped phenotype of the mutants is due to elongation of the proximal portion of the tubule (**Supplementary Fig S5; Supplementary Videos S1,S2**). This result phenocopies the looped appearance of the elongated proximal tubule in the macrobasic mastigophores found in other sea anemones (**Fig 3H**), suggesting a parallel regulation scheme may control tubule morphogenesis in each lineage of cnidocytes. Thus, *NvSox2* has different roles in the development of small and large basitrichous isorhizas, confirming these are distinct cell types and not just two size classes of a single cell type.

Next, we examined the development of the native (gracile) spirocytes (**Fig 3I**) to determine if *NvSox2* knockout affected specification of this cell type. We have identified the first positive and specific marker of gracile spirocytes in *N. vectensis*, a Ca^2+^-binding protein called Calumenin F (*Caluf*) (*29*). *Caluf* is coexpressed in cells labeled with anti-Mcol4 antibody in the developing tentacles of WT polyps but is never co-expressed with nematocyte-specific gene *Mcol1* (**Fig 3J**) (*25*). Knockout of *NvSox2* did not affect the timing or the distribution of cells expressing *Caluf* (**Fig 3K**; **Supplementary Fig S4**) and TEM confirmed the morphology of the gracile spirocytes from WT animals did not differ from that of *NvSox2* mutants (**Fig 3L**). Considering there was also no effect of *NvSox2* knockout on the nematocytes that populate the internal digestive tissues (microbasic p-mastigophores; **Supplementary Fig S6**), these data confirm the role of *NvSox2* in driving cell type specification is restricted only to the small basitrichous isorhiza lineage of nematocytes in *N. vectensis*.

### NvSox2 is present in only a subset of cnidarians

*Sox* genes are ubiquitous among animal genomes and cluster into 8 major groups (*SoxA-SoxH*) (*15*), at least four of which were likely present in the common ancestor of cnidarians and vertebrates (B,C,E,F). Previous phylogenetic analyses of *Sox* genes from *N. vectensis* have suggested that *NvSox2* is either a cnidarian-specific ortholog (*16*) or a member of the *SoxB* clade (*20, 30, 31*). To test these hypotheses further, we built a maximum likelihood phylogeny of *Sox* genes from 15 cnidarians and 6 bilaterians (**Fig 4A; Supplemental Data S1**). With this extensive taxon sampling, we find strong support for the monophyly of a clade including *NvSox2* orthologs from sea anemones and corals but weak support linking the *NvSox2* clade to any known bilaterian *Sox* genes (**Fig 4B,C**). We find no evidence of an *NvSox2* ortholog in medusozoans, octocorals, or cerianthids (tube anemones). This suggests *NvSox2* arose in the stem ancestor of non-cerianthid hexacorals, coincident with the origin of robust spirocytes.

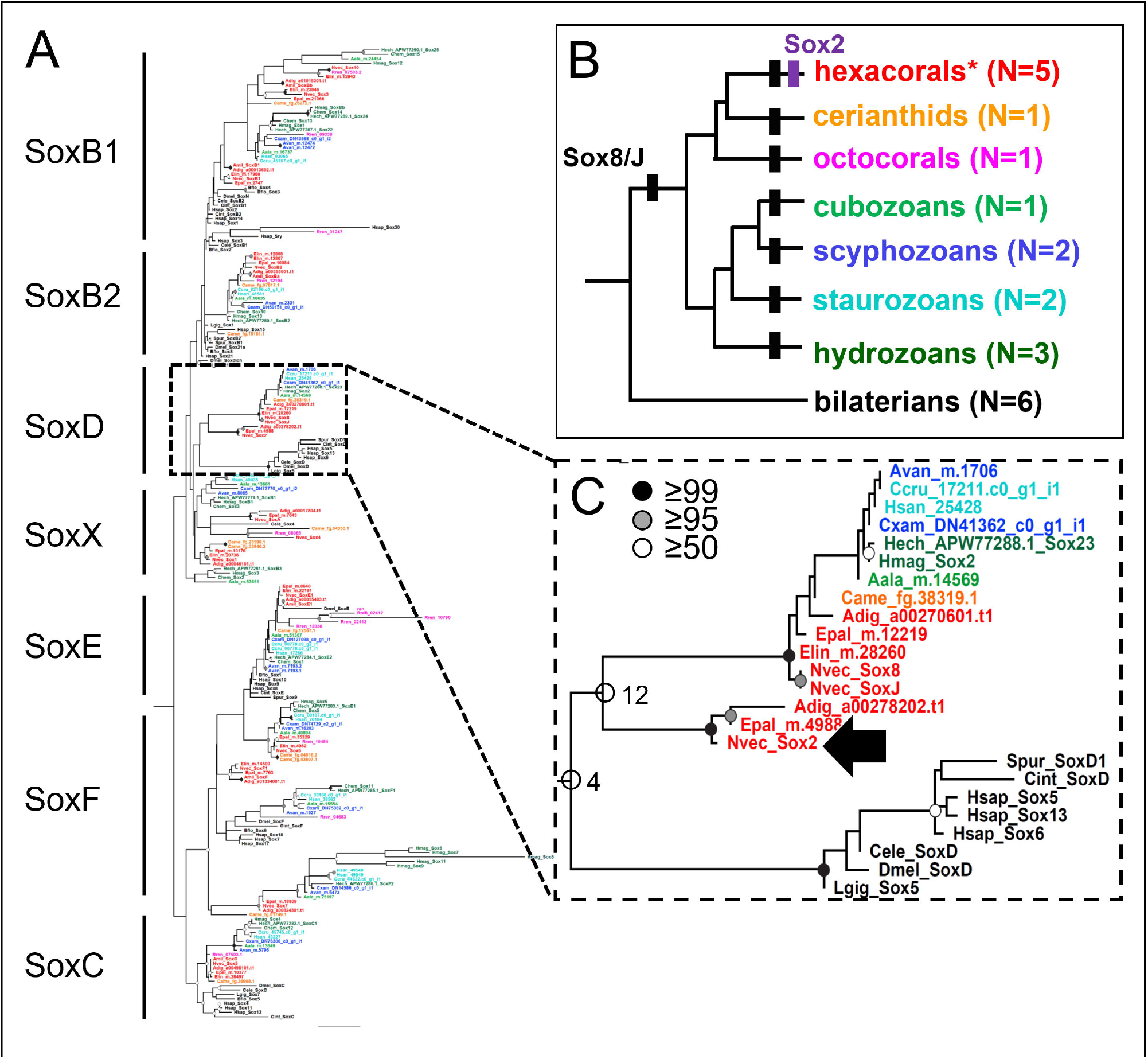

### Cell type homeosis explains cnidocyte diversity

The data presented here suggest that “nematocyte” and “spirocyte” are not phylogenetically distinct cell types but instead represent two alternative fates within a single cell lineage (**Fig 5A-C**). Further, while small and large basitrichous isorhizas have been assumed to be two size classes of the same cell type, we show that these are two molecularly distinct types of nematocyte, as only the former took on a spirocyte fate in mutant animals (**Fig 5D**). Indeed, homeotic control of cell identity could explain how the cnidom of two closely related cnidarians could differ quite dramatically (**Fig 5E,F**). Burrowing sea anemones are found in a wide variety of habitats and provide a valuable illustration of this phenomenon: while some species are found in muddy/sandy habitat (like *Nematostella vectensis* and *Edwardsiella ignota*), others burrow into sea ice (like *Edwardsiella andrillae*)(*32*). An evolutionary transition between the small basitrichous isorhizas found in the body wall of *E. ignota* and the robust spirocytes in the body wall of *E. andrillae* could be explained by downregulation or loss of an *NvSox2* ortholog in the latter. Likewise, homeotic control of developmental plasticity could also explain the transition from microbasic b-mastigophores to isorhizas in the body wall of *Edwardsiella lineata* and *E. carnea* as these sea anemones transition from a free-living state to a parasitic stage spent in the digestive tract of ctenophores (*33, 34*). A possible loss of the *NvSox2* ortholog in *E. lineata* (**Fig 4**) suggests a similar single-gene regulatory logic may also control binary fate decisions between other cnidocyte subtypes, though the identity of the regulatory gene remains elusive (**Fig 5D-F**).

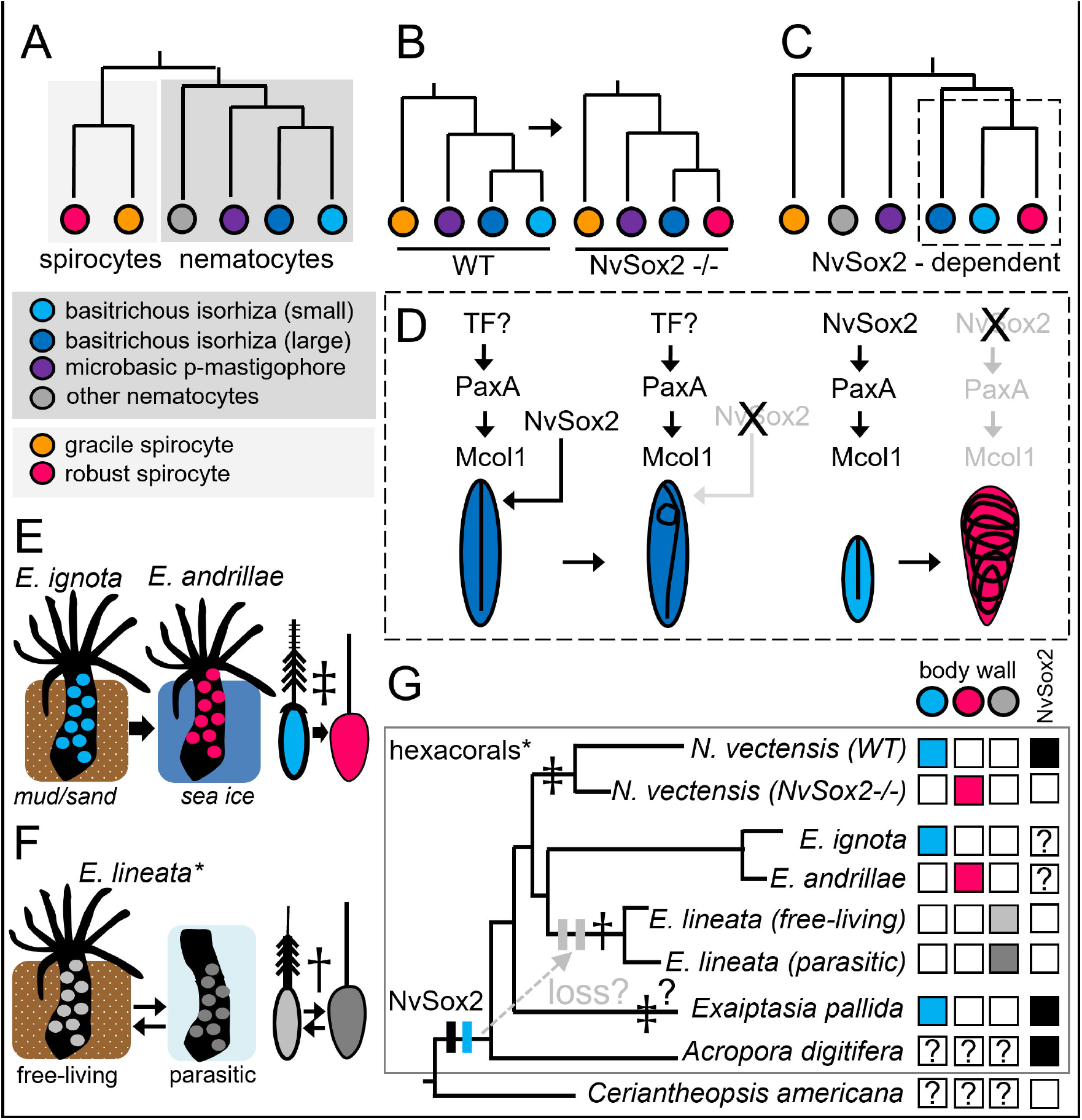

## Conclusions

We have shown that knockout of a single transcription factor induced nematocytes to take on the morphology (**Fig 2**) and molecular signature (**Fig 3**) of spirocytes. Importantly, the mutant spirocytes also retain the ability to discharge, confirming this mutation is functionally relevant and is, therefore, a *bona fide* homeotic transformation (*7*). This is the first evidence of homeotic control of cell fate in a cnidarian, suggesting that cellular homeosis is an ancient mechanism that contributed to the expansion of biodiversity that emanated from the last common ancestor of cnidarians and bilaterians, over 800MYA (*35*). Specifically, our ability to resurrect an ancestral cell type by manipulating a single gene demonstrates how homeosis links developmental flexibility with environmental selection pressures to drive the emergence of novel cell types. In support of this, a single regulatory gene was found to control a homeotic shift in butterfly wing color pattern, demonstrating a mechanism by which rapid evolution can occur when a complex phenotype is controlled by a single switch (*36*). The ability to silence ancestral cell types while retaining the instructions for their specification allows animals extreme flexibility to explore new habitats. The ability to rapidly switch between multiple unique cell identities with a single gene may well be the most efficient way for animals to adapt to local conditions, suggesting this phenomenon may be widespread among animals.

The observation that NvSox2 controls tubule morphogenesis – not specification – in a second lineage of nematocyte (large basitrichous isorhizas), recapitulates phenotypic variation in another lineage of cnidocytes (mastigophores). These results uncover a mechanism for generating the unique identities of each cnidocyte subtype by varying the combinatorial expression of genes controlling individual modules of the phenotype, similar to the modular control of neuronal identity described from nematodes (*37*). Specifically, the fact that *NvSox2* controls only the morphology of the proximal tubule suggests it should be possible to find individual genes that similarly control the morphology of the distal tubule, the composition of the capsule wall, and the nature of the apical structures. This “modular phenotype” framework would explain how evolution could mix and match genes controlling each aspect of the phenotype of these organelles to produce the diverse array of cnidocyte subtypes present today.

## Acknowledgements

Electron microscopy was performed at the Biological Electron Microscope Facility (BEMF) at the University of Hawai’i, Mānoa, and at the Center for Electron Microscopy and Analysis (CEMAS), the Campus Microscopy and Imaging Facility (CMIF), and the OSU Comprehensive Cancer Center (OSUCCC) Microscopy Shared Resource (MSR) at The Ohio State University (OSU). We are grateful to Jeffrey Tonniges for his assistance with sample preparation for TEM at OSU.

## Author Contributions

LSB, CE, BMF, FH, and MQM performed laboratory procedures including development of CRISPR reagents, microinjection, in situ hybridization, cloning and sequence analysis, and confocal microscopy. LSB, AJR, and MD performed electron microscopy and analyzed mutant cell phenotypes. LSB and JFR performed phylogenetic analyses. LSB conceived of the study and wrote the manuscript. All authors edited the manuscript and approved the final draft.

## Data availability

All supporting data and code are provided in the supplement to this manuscript.

## Brief Methods

The NvSox2 locus was deleted using CRISPR/Cas9 genome editing as previously described (*38, 39*). Briefly, five guide RNAs targeting different regions of the locus were injected together with Cas9 protein and a fluorescent dextran into fertilized eggs. Animals were raised to reproductive age, crossed with WT animals, and genome editing was confirmed in their progeny by genomic DNA from tentacle clips. One mutant male and one mutant female were used to generate the F2 generation, in which gene expression and other cell/tissue analyses were performed. Gene expression was inhibited using morpholinos as previously described (*28*). qPCR and FISH were also performed as described (*28*). A method for labeling nematocyte tubules using H_2_O_2_-Cy3 tyramide was developed for this study. Briefly, primary polyps (4-tentacle stage) were fixed as described previously for in situ hybridization. After extensive rinsing in PBS with 0.1% Tween, polyps were incubated in Cy3-tyramide (1:500 in PBS) for 20 minutes and Cy3-tyramide with 0.1% H_2_O_2_ for 45 mins. Polyps were then rinsed in PBS/Tween to remove excess Cy3-tyramide, counterstained with DAPI, mounted in glycerol, and imaged on a Zeiss 710 confocal microscope. To induce discharge of cnidocytes, polyps were immobilized for 5 mins in 7.14% MgCl_2_ and mounted under coverslips with clay feet. An infrared laser ablation system (XYClone, Hamilton Thorne) was used to induce discharge from tentacle tip cnidocytes with 100% power and a 500us pulse. Phylogenetic analysis of HMG domains was performed using translated transcriptomes as described previously (*40*). Full methods are provided in the Supplementary Information.

Kayal et al 2018(*41*)

