## Supplementary Figure S5 for "Knockout of a single *Sox* gene resurrects an ancestral cell type in the sea anemone *Nematostella vectensis*"

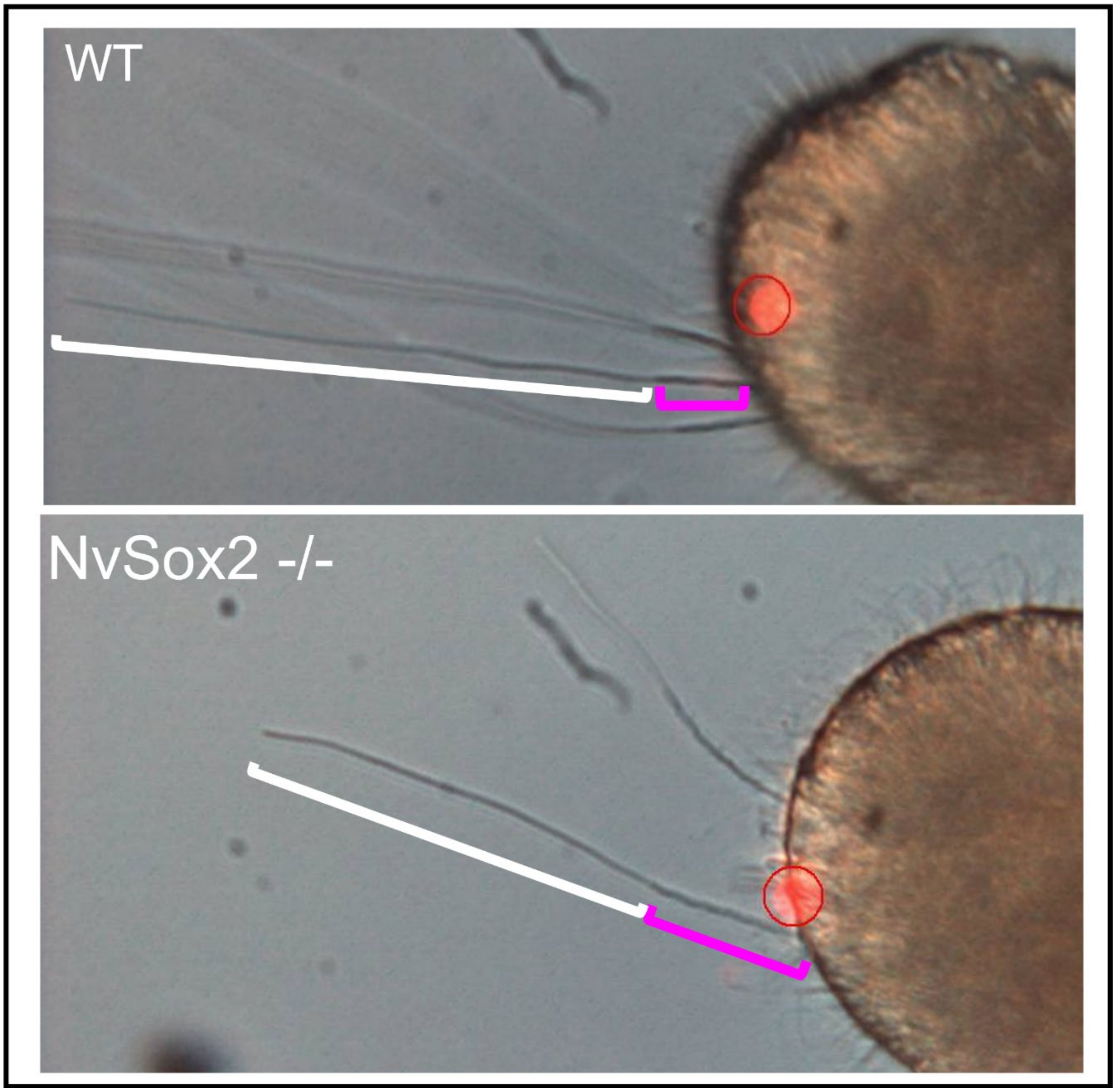


**Fig. S5**. Still images captured from videos of laser-induced cnidocyte discharge. Magenta brackets indicated the proximal (spiny) portion of the discharged tubule from a large basitrichous isorhiza, which is longer in *NvSox2* mutants than WT polyps. Red circled area shows the laser target (moved from target area after discharge).
