## Supplementary Figure S6 for "Knockout of a single *Sox* gene resurrects an ancestral cell type in the sea anemone *Nematostella vectensis*"

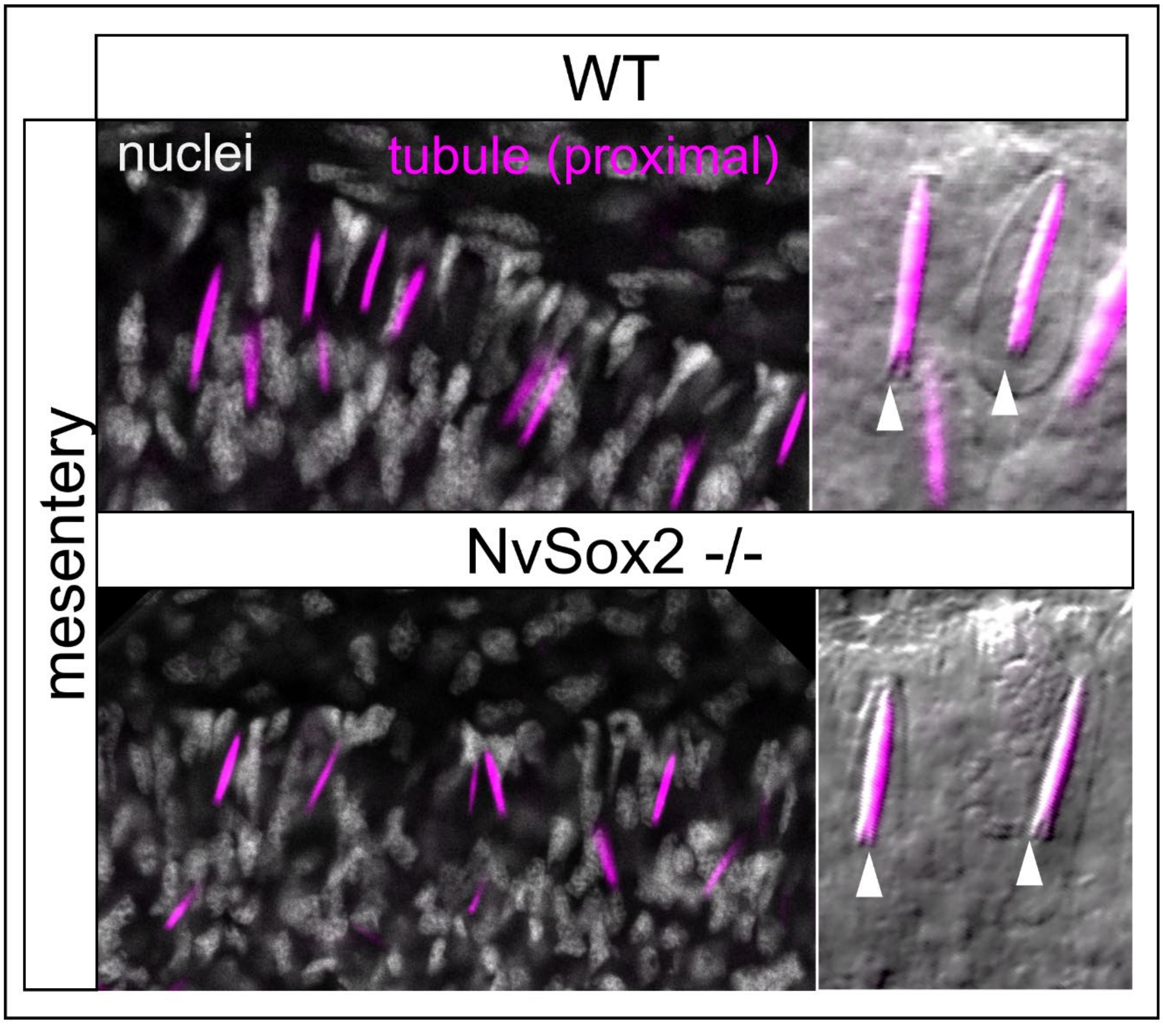


**Fig. S6**. The morphology of microbasic p-mastigophores in the mesenteries of WT and *NvSox2* mutant primary polyps did not differ (arrowheads in inset point to v-notch unique to this cell type).
