## Supplementary Table S1 for "Knockout of a single *Sox* gene resurrects an ancestral cell type in the sea anemone *Nematostella vectensis*"

**Table S1.** Sequences for single guide RNAs (sgRNAs) and genotyping primers for *NvSox2* mutants.

| **name** | **sequence (5'->3')** | **notes** |
|---|---|---|
| NvSox2_gRNA1 | TAGCGGGCGATCTATTCATTTGG | guide RNA (5'UTR) |
| NvSox2_gRNA2 | CCATGAACGCGTATATGGTATGG | guide RNA (coding sequence) |
| NvSox2_gRNA3 | CGAGTGGAATTCGCTCACTTTGG | guide RNA (coding sequence) |
| NvSox2_gRNA4 | CCAACCAATGGCTTCCCATATGG | guide RNA (coding sequence) |
| NvSox2_gRNA5 | CACCCATAGACGCACGAAGTGGG | guide RNA (coding sequence) |
| NvSox2_Cr_F primer | GGATAAGACATTCCACCCGTTTTC | forward primer |
| NvSox2_Cr_R primer | CGTTCGGTTTCTGTGGTAAAACTC | reverse primer |
