## Supplementary Information for "Knockout of a single *Sox* gene resurrects an ancestral cell type in the sea anemone *Nematostella vectensis*"

**This PDF file includes:**

Materials and Methods
Captions for Movie S1, Movie S2, Data S1, and Data S2

**Other Supplementary Materials for this manuscript include the following:**

Data S1: Alignment for maximum likelihood tree (.txt).
Data S2: Survey of robust spirocytes in other cnidarians (.xls).

### Materials and Methods

**Genome Editing**: The NvSox2 locus was deleted using a modification of two CRISPR/Cas9 protocols published previously (Ikmi et al 2014, Servetnick et al 2017). Briefly, five guide RNAs were designed to target different sites in the NvSox2 locus, four in the coding sequence and one in the 5' UTR (Supplementary Fig. S1, Supplementary Table S1). Guide RNAs were purchased from Synthego (USA) and reconstituted to 30uM in nuclease-free water (Synthego, USA) following the manufacturer's instructions. Cas9 protein was purchased from PNABio (USA) and reconstituted to 1mg/ml in nuclease free water (Ambion, USA) following the manufacturer's instructions. Guide RNAs were mixed together with Cas9 in a ratio of 5:4 (vol:vol) gRNA:Cas9, incubated at room temperature for 10 minutes, and injected into zygotes with 0.2 mg/ml Alexa-555 RNAse free dextran (Invitrogen D34679) and nuclease-free water (Ambion AM9937). To determine if there were non-specific effects of guide RNAs or Cas9, control embryos were injected with guide RNAs but no Cas9 protein or Cas9 without guide RNAs. These animals developed normally and were not examined further. Mosaic F0 NvSox2 mutant embryos were raised to reproductive maturity at 16C and spawned to generate the F1 generation. Genomic DNA was extracted from tentacle tips of F1 polyps and sequenced to check for mutations. One F1 female and one F1 male were identified to have significant deletions in the NvSox2 locus (Supplementary Fig. S1) and were used to produce the F2 generation. All further cell/tissue analyses were performed in the F2 generation.

**Cell and tissue analysis:** To assay effects of NvSox2 knockout on cnidocyte specification, developing cnidocytes were labeled, imaged, and counted as described previously (Babonis and Martindale, 2017) using an antibody directed against minicollagen4 (Mcol4) (Zenkert et al 2011*)*. Embryonic expression of NvSox2 and several known cnidocyte genes was examined using in situ hybridization (Babonis and Martindale, 2017). To determine if NvSox2 was part of the cnidocyte gene regulatory network, we knocked down SoxB2 and PaxA using published morpholinos and methods that have been described previously (Babonis and Martindale, 2017). For qPCR analysis, five replicates of each condition (WT, control MO, SoxB2 MO, and PaxA MO) representing five independent injections performed on different days were compared using the delta-delta CT method and the PCR package (Ahmed and Kim, 2018) in the R statistical computing environment (R Core Development Team). Expression values in Fig. 2B are presented as fold-change, relative to expression in uninjected embryos and statistical significance was calculated from the comparison of SoxB2 MO and PaxA MO-injected embryos relative to control MO-injected embryos. Significant differences (where $p<1E-02$) are indicated by different letters in all figures representing quantitative comparisons.

**Electron Microscopy (Fig. 1,2,3):** Samples were prepared for TEM and SEM analysis as described previously (in Reft and Daly, 2012 and Dubuc et al., 2014). Briefly, for TEM, juvenile polyps were either fixed in 2.5% glutaraldehyde (in phosphate buffer) and postfixed in 1% $OsO_4$ (in phosphate buffer) or fixed in 4% glutaraldehyde (in cacodylate buffer) and postfixed in 2% OsO4 (in cacodylate buffer). Specimens were then dehydrated in a graded ethanol series and embedded in epoxy resin. Thin sections (60 -70nm) were stained with 2% or 5% aqueous uranyl acetate for 15min and then for 5 min in Reynolds lead citrate. Micrographs were taken on a Technai B2 Spirit TEM at The Ohio State University, Columbus OH, or a Hitachi HT7700 TEM at the University of Hawai'i (UH), Mānoa. In preparation for SEM, some samples were treated with 1M Sodium Citrate for 15 mins to induce cnidocyte discharge as previously described (Reft and Daly, 2012). These samples were rinsed in DI water and fixed in 70% ethanol before being dehydrated through a graded ethanol series. Some samples discharged fortuitously and were fixed as for TEM in 4% glutaraldehyde followed by 2% $OsO_4$ before being dehydrated in ethanol. All samples were critical point dried, sputter-coated, and imaged on either a FEI Quanta 200 scanning electron microscope at OSU or a Hitachi S-4800-I field-emission scanning electron microscope at UH.

**Analysis of Mcol4 and Mcol1 expression (Fig. 3C)**: Developing animals were fixed for examination at the tentacle bud stage (240 hours post fertilization). We performed a combination of in situ hybridization and immunohistochemistry on the same tissues as previously described (Babonis and Martindale, 2017). Labeled tissues were then mounted in glycerol on glass slides, imaged with a Zeiss 710 confocal microscope, and z-stacks were rendered into 3D images using Imaris software (Oxford Instruments, USA). Regions of interest were demarcated using the crop tool in Imaris and labeled cells were counted by eye in a 100x100um square region of the animals immediately aboral to the tentacle buds. Data are presented as percent of cells co-labeled with Mcol1 and Mcol4 (magenta bars) relative to the total number of Mcol4-labeled cells (aqua bars) adjusted to 100% in WT. Error bars represent standard deviation for N = 10 tentacle bud stage animals examined in each treatment. We used a Mann Whitney U test to compare the number of cells of each type in WT and NvSox2 mutants and found no significant difference in the total number of Mcol4-labeled cells ($p = 0.623176$) and a significant decrease in the number of cells co-labeled with Mcol4 and Mcol1 ($p=0.000157$) in NvSox2 mutants.

**Squash preps (Fig. 3E,F):** Embryos were raised at 16C until they metamorphosed into primary polyps (~14 days post fertilization). Polyps were immobilized in 7% $MgCl_2$, fixed for 1 min in 4% PFA/glut and for 2h in 4% PFA at 4C. Tissues were then washed several times in PTw to remove fixative, mounted on a glass slides in minimal 80% glycerol, and squashed/smeared under a coverslip to dissociate the tissues into individual cells. Coverslips were permanently mounted with nail polish. Nematocytes and mutant cell types were counted by eye using the 40X objective on a Zeiss Axioscope M2. A single polyp was mounted under each coverslip and N = 12 total polyps per condition (wildtype or mutant) were examined. We made three, non-overlapping transects through the squashed tissue counting every cell within the field of view for each transect. We then added the total number of robust spirocytes and nematocytes from all three transects on a single animal to estimate the total proportion of cnidocytes that were robust spirocytes in WT and Sox2 mutants. The abundance of mutant cell types was low (<1%) in WT animals and increased significantly to nearly 50% in mutant animals (Mann Whitney U; $p = 0.000032$).

**Nematocyst tubule analysis with $H_2O_2$ (Fig. 3H):** Recently metamorphosed polyps (~14 days post fertilization) were fixed for 1min at 25C in PFA/glut and for 1h at 4C in 4% PFA, as for in situ hybridization. Fixed polyps were washed twice in DI water to remove PTw and stored in 100% methanol at -20 until use. For staining, tissues were rehydrated from methanol into PTw at 25C, treated for 20 mins with 0.1% $H_2O_2$ and then incubated in 1% tyramide-Cy3 (in 0.1% $H_2O_2$) for 45 mins in the dark. Excess tyramide/ $H_2O_2$ was removed with three washes in PTw and polyps were counterstained with 1uM DAPI for 30 mins at 25C before being mounted on glass slides in glycerol and compressed under a coverslip with clay feet. Imaging was performed the same day as labeling on a Zeiss 710 confocal. Labeled tubules from N = 11 polyps per condition (wildtype or mutant) were counted by eye and nuclei (labeled with DAPI) were counted automatically using the Spots tool in Imaris. The abundance of nematocytes (#tubules / #nuclei) was compared between treatments with a Mann Whitney U test. The total number of nematocyst tubules decreased significantly in NvSox2 mutants, relative to WT polyps ($p = 0.000160$).

**Maximum Likelihood phylogeny (Fig 4):** Before starting any analyses, experiments were planned and described in a phylotocol (Ryan and Debiasse, 2018). Subsequent modifications to the analyses were noted and justified in the phylotocol. Many genes besides *Sox* genes include an HMG box, so searching for *Sox* genes using just the HMG hidden markov model (HMM) produced many non-target sequences. To identify Sox genes specifically, we generated a custom HMG HMM from a published *Sox* gene alignment (Schnitzler et al, 2014) after removing the outgroup sequences (Tcf/Lef and Capicua/CIC) using hmmbuild (hmmer.org). We then used this custom HMM to search for *Sox* genes in translated transcriptomes from 15 cnidarians and six bilaterians. The abbreviations for the cnidarian taxa we used are as follows: **Aala** - *Alatina alata*, **Adig** – *Acropora digitifera*, **Amil** - *Acropora millepora*, **Epal** - *Exaiptasia pallida*, **Avan** - *Atolla vanhoeffeni*, **Ccrux** - *Calvadosia cruxmelitensis*, **Came** – *Ceriantheopsis americana*, **Chem** – *Clytia hemisphaerica*, **Cxam** – *Cassiopea xamachana*, **Elin** – *Edwardsiella lineata*, **Hech** – *Hydractinia echinata*, **Hmag** – *Hydra magnipapillata*, **Hsan** – *Haliclystus sanjuanensis*, **Nvec** – *Nematostella vectensis*, **Rren** – *Renilla reniformis*; and for bilaterians: **Bflo** – *Branchiostoma floridae*, **Cint** – *Ciona intestinalis*, **Cele** – *Caenorhabditis elegans*, **Dmel** – *Drosophila melanogaster*, **Hsap** – *Homo sapiens*, **Lgig** – *Lottia gigantea*, **Spur** - *Strongylocentrotus purpuratus*. We used this custom HMM in combination with our hmm2aln script https://github.com/josephryan) to generate an alignment that included the original sequences used to generate the HMM. We then removed all ctenophore, sponge, and placozoan sequences from this alignment and generated trees.

The full methods for our phylogenetic analysis have been described previously (Babonis et al 2019). Briefly, we used the model finder feature with IQ-TREE to identify the best substitution model for the alignment. We then performed three maximum likelihood analyses, in parallel, using: RAxML with 25 maximum parsimony starting trees, RAxML with 25 random starting trees, and a default run with IQ-TREE. We then compared maximum likelihood values from the outputs of all three analyses to select the best tree and performed 1000 rapid bootstraps using RAxML for branch support. The final tree file was modified in FigTree and Adobe Illustrator for presentation.

**Movie S1.**

Discharge of nematocytes from the tentacle tips of WT polyps using an infrared laser ablation system.

**Movie S2.**

Discharge of nematocytes from the tentacle tips of NvSox2 mutant polyps using an infrared laser ablation system.

**Data S1. (separate file)**

Alignment for maximum likelihood tree (.txt).

**Data S2. (separate file)**

Literature survey of reports of robust spirocytes in other cnidarians (.xls)
