## Supplementary figures and images for "Knockout of a single *Sox* gene resurrects an ancestral cell type in the sea anemone *Nematostella vectensis*"

### Supplementary Figure S1

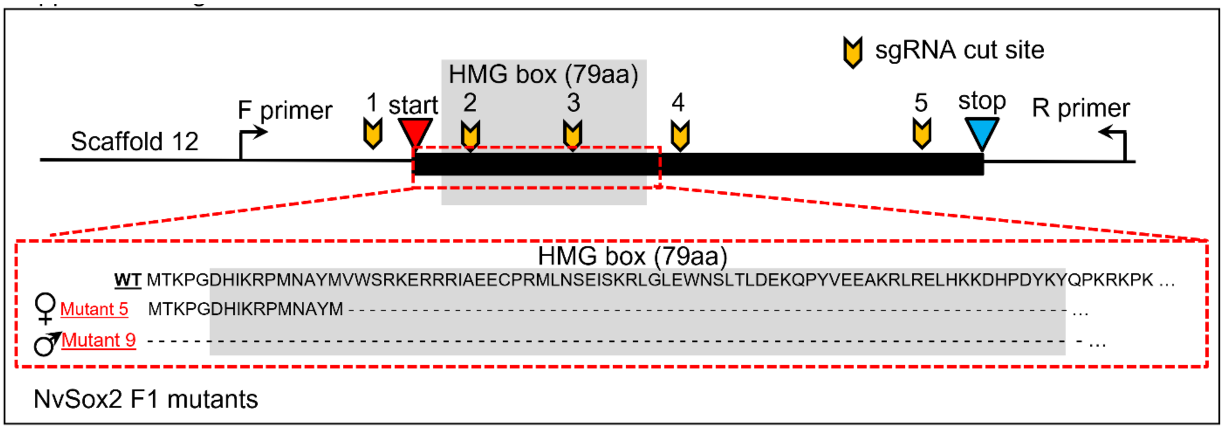


**Fig. S1**. Genomic locus of *NvSox2* with guide RNA (sgRNA) cut sites and F1 CRISPR mutations indicated.
