## Supplementary Figure S2 for "Knockout of a single *Sox* gene resurrects an ancestral cell type in the sea anemone *Nematostella vectensis*"

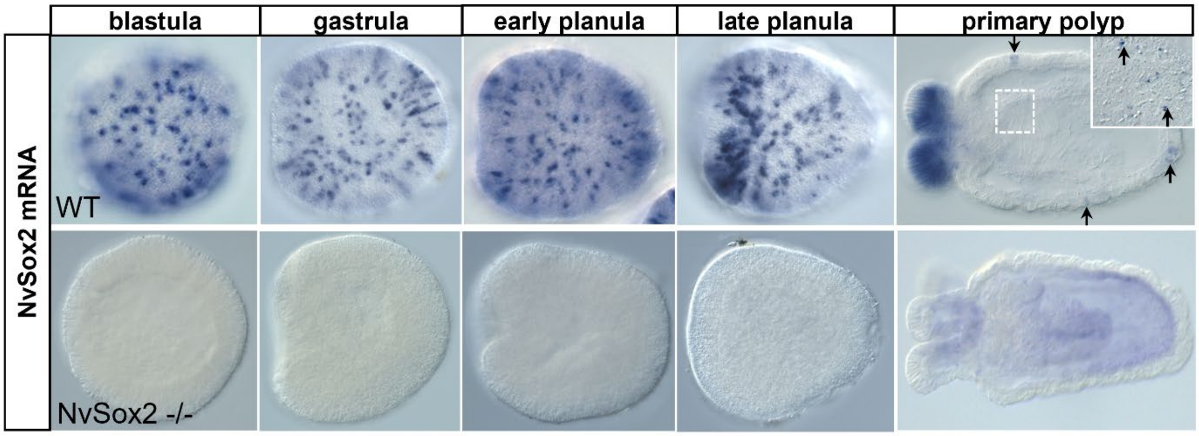


**Fig. S2**. Expression profile (in situ hybridization) of *NvSox2* in WT and *NvSox2* mutant embryos showing complete loss of NvSox2 in mutants.
