## Supplemental Figure S3 for "Knockout of a single *Sox* gene resurrects an ancestral cell type in the sea anemone *Nematostella vectensis*"

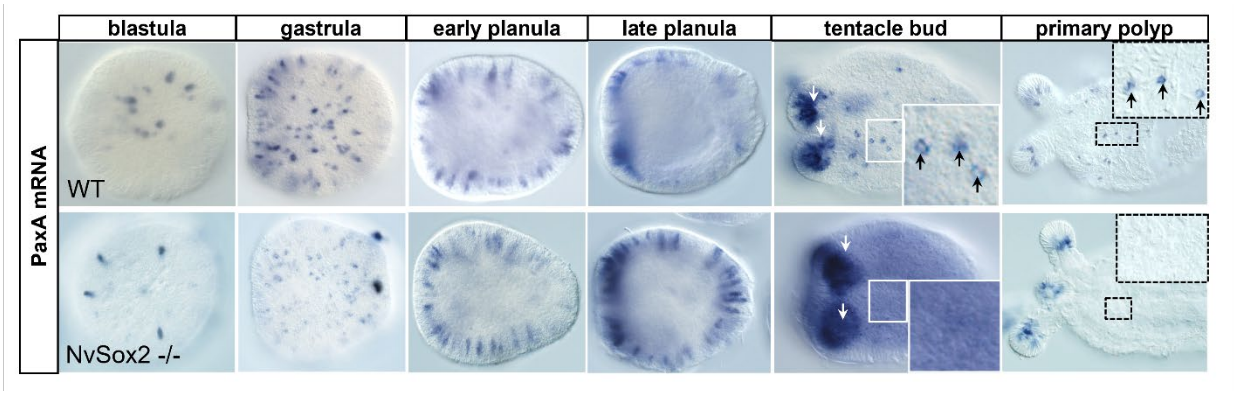


**Fig. S3**. Expression profile (in situ hybridization) of *PaxA* in WT and *NvSox2* mutant embryos showing loss of *PaxA* only in the body wall of mutants.
