## Supplemental Figure S4 for "Knockout of a single *Sox* gene resurrects an ancestral cell type in the sea anemone *Nematostella vectensis*"

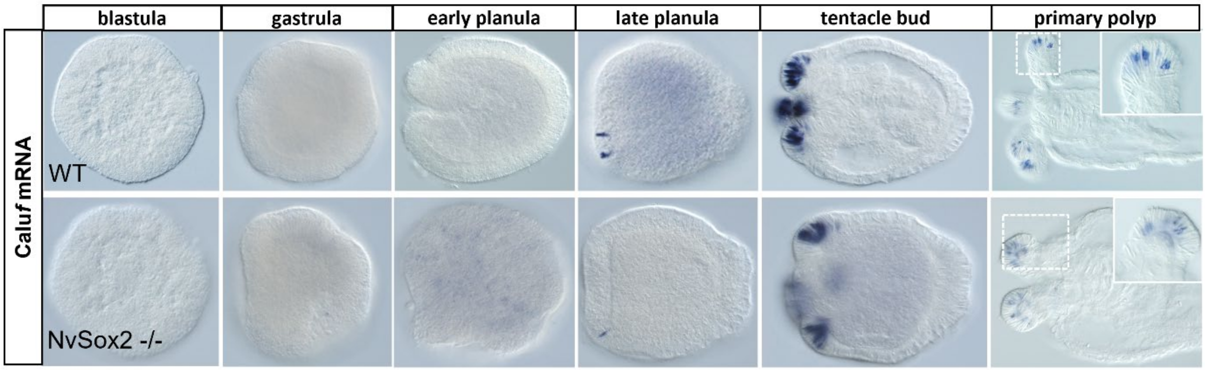


**Fig. S4**. Expression profile (in situ hybridization) of *Caluf* in WT and *NvSox2* mutant embryos showing no effect of *NvSox2* knockout on *Caluf* expression.
